# Patient-derived Brain Organoids Reveal Divergent Neuronal Activity Across Subpopulations of Autism Spectrum Disorder

**DOI:** 10.1101/2025.04.27.650821

**Authors:** Nisim Perets, Liya Kerem, Nir Waiskopf, Noa Horesh, Itay Goldman, Jasmine Avichzer, William Tobelaim, Milcah Barashi, Liat David, Ariel Tenenbaum

## Abstract

Patient-derived brain organoids have emerged as a powerful model for investigating the mechanisms underlying neurological and psychiatric disorders. They provide novel insights into autism spectrum disorder (ASD), a heterogeneous neurodevelopmental condition whose underlying mechanisms remain poorly understood. Recent advancements in generating electrophysiological functional 3D brain organoids enable the study of molecular and network-level neuronal activity.

Here, we aimed to characterize the neurophysiological underpinnings of ASD by comparing electrophysiological properties of brain organoids derived from eleven individuals diagnosed with autism spectrum disorder -10 with monogenic syndromic ASD across five genetic subtypes, and 1 with idiopathic ASD -to organoids derived from 4 neurotypical control individuals. We identified distinct differences in baseline activity (resting state) and evoked responses (synaptic plasticity and network dynamics) across ASD subgroups. To comprehensively assess these differences, we applied dimensionality reduction and machine learning (principal component analysis, PCA) to integrate multiple electrophysiological features into a unified framework.

Our findings reveal subtype-specific neurophysiological alterations in ASD brain organoids, offering mechanistic insights into ASD heterogeneity and potential applications for early diagnostics, drug screening, and therapeutic development.

## Introduction

The prevalence of neurological and neuropsychiatric disorders worldwide is striking. According to epidemiological studies from 2021, more than 3 billion people worldwide live with one or more neurological or neuropsychiatric conditions, affecting approximately 43.1% (40.5-45.9%) of the global population^1^. Autism Spectrum Disorders (ASD) are neurodevelopmental conditions characterized by persistent deficits in social communication and interaction, along with restricted repetitive behaviors, with symptoms varying widely in severity^2^. Worldwide, ASD affects approximately 75 million individuals, ranking 12th among all neurological and psychiatric conditions and 5th among neuropsychiatric disorders in terms of prevalence^3^.

ASD is driven by a complex interplay of genetic, biological, and environmental factors that disrupt early brain development^4^. Factors such as prenatal exposure to infections, maternal immune activation, and metabolic or environmental insults may also contribute, usually leading to idiopathic ASD.^5,6^ Yet, in recent years, hundreds of genes have been implicated in ASD risk, classified as syndromic ASD.

Syndromic ASD, stemming from single-gene mutations, offers valuable insights into the disorder’s biological basis. Notable examples include SHANK3 haploinsufficiency syndrome, PPP2R5D-related disorder (Jordan’s syndrome), SCN2A-related disorder, GRIN2B-related neurodevelopmental syndrome, and STXBP1-associated disorders, all of which impact synaptic function, neural excitability, and circuit development^7,8^. The genetic mutations driving these syndromes disrupt essential molecular pathways responsible for neuronal network development and function. **SHANK3 haploinsufficiency**, seen in Phelan-McDermid syndrome, leads to impaired synaptic scaffolding at excitatory synapses, disrupting glutamatergic signaling and synaptic plasticity^9,10^. **PPP2R5D mutations** (Jordan’s syndrome) alter the regulatory subunit of protein phosphatase 2A, impacting neuronal development and signal transduction, particularly pathways involved in brain growth and cytoskeletal regulation^11^. **GRIN2B-related syndrome**, caused by mutations in the gene encoding the GluN2B subunit of the NMDA receptor, results in defective excitatory neurotransmission and altered calcium signaling, impairing learning-related plasticity^12^. Mutations in **SCN2A**, which encodes a voltage-gated sodium channel, affect neuronal excitability and are strongly associated with both ASD and epilepsy due to disrupted action potential generation^13^. Lastly, **STXBP1 mutations** impair presynaptic vesicle release by disrupting the SNARE complex, leading to widespread deficits in neurotransmitter release and cortical network function^14,15^. Together, these mutations converge on different pathways affecting synaptic transmission, plasticity, and network connectivity, providing critical insights into the molecular underpinnings of ASD. Though it captures about 10% of the overall ASD population, it can hold an important key to unlock impaired mechanisms of idiopathic populations^16^.

However, despite the growing abilities in mapping new syndromic populations, the extrapolation from genes to neuronal activity patterns is still challenging, especially in human patients. This is compounded by the limited availability of viable human patient-derived brain tissue with preserved functional integrity. Furthermore, traditional animal models often fail to fully capture human-specific features of ASD, limiting the translational relevance of their findings ^20^.

Brain organoids - three-dimensional, self-organizing structures derived from induced pluripotent stem cells (iPSCs) ^21^, recapitulate key features of early brain development, including the emergence of distinct brain regions and cell types. Crucially, they retain the patient’s specific genetic background and molecular fingerprint^22–24^, enabling the modelling of disease-relevant phenotypes in a human context. As such, patient-derived brain organoids have emerged as a cutting-edge in vitro system, offering valuable molecular insights into the mechanisms underlying ASD and other neurological and neuropsychiatric disorders^25–27^.

Notably, they can also replicate key aspects of neuronal diversity and network connectivity, providing a valuable platform to study synaptic plasticity and disrupted neural circuitry that can unlock new mechanisms underlying neuropsychiatric disorders^28^.

Here, we analyzed 18 electrophysiological features of patient-derived brain organoids and used principal component analysis (PCA) to reduce them into a 3D space for comparison. Organoids from the same individual showed minimal intra-subject variability, and control patients clustered tightly, reflecting low inter-subject variability. In contrast, syndromic and idiopathic ASD samples were widely dispersed, highlighting significant heterogeneity. Interestingly, even patients with the same genetic mutation in the syndrome displayed distinct electrophysiological patterns, placing them far apart in the 3D space.

## Materials and Methods

### Participant recruitment and sample collection

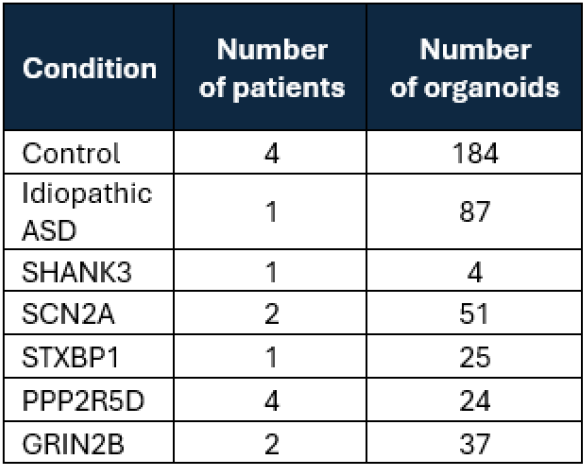

The differences in the number of brain organoids among the various groups result primarily from challenges in forming them across different lines while aiming to maintain minimal protocol interventions. A clinical assessment of the patients and their phenotype is summarized in Table S1.

* The idiopathic ASD and one of the control lines are a father and son. The rest of the patients have no familial relationship.

### Urine collection and iPSC generation

iPSC lines were reprogrammed from urine-derived epithelial cells or acquired from the Simons Foundation Autism Research Initiative (SFARI). Urine samples were collected from healthy volunteers and patients with ASD following approval by the Institutional Review Board (IRB) and the ethics committee of Hadassah Medical Organization (Research application: HMO-0021-22), and conducted according to institutional ethics committee guidelines. Epithelial cells were precipitated from urine by centrifugation at 400g for 10 min and later selected and expanded on plates coated with iMatrix-511 (T304, Takara-Clontech), and media containing DMEM/Ham’s F12 1:1, Lonza™ REGM™ Renal Epithelial Cell Growth Medium SingleQuots™ Kit (CC-4127), human serum (AC-001-1B, Access Cell Culture) and PSA as antibiotics^29,30^. Subsequently the urine-derived epithelial cells were reprogrammed using Epi5™ Episomal iPSC Reprogramming Kit (A15960, Thermo Fisher Scientific^31^), and kept in liquid nitrogen till organoid generation.

iPSC lines acquired from SFARI: 15653-x1, 16020-x1, 15655-x1, 16051-x1, 15413-x1, 15265-x1, 15473-x1, 15936-x1, 15912-x1.

### iPSC immunostaining

Immunostaining was performed to analyze the expression of key pluripotency markers in induced pluripotent stem cells (iPSCs). The cells were cultured under standard conditions until they reached 40-50% confluency. Following this, iPSCs were fixed with 4% paraformaldehyde for 20 minutes at room temperature (R.T.). Permeabilization was achieved using Triton X-100 for 45 minutes at R.T. After blocking with bovine serum albumin for 1 hour, the cells were incubated overnight at 4°C with primary antibodies: Mouse anti-Sox2 (SC-365823) and Anti-Oct-3/4 Antibody (SC-5279) from Santa Cruz Biotechnology, and NANOG Monoclonal Antibody (eBioMLC-51, cat. 14-5761-80, Invitrogen) at concentrations recommended by the manufacturer. The expression of Sox2 and Oct4 was analyzed as markers of pluripotency (data not shown), as both are critical for maintaining stem cell self-renewal and undifferentiated states. NANOG staining was included to further confirm pluripotency, as it is essential for the maintenance of iPSC self-renewal and inhibiting differentiation. Following incubation with primary antibodies, the cells were washed with PBS and incubated with secondary antibodies conjugated to fluorophores from Invitrogen: Goat anti-Mouse IgG (H+L) Alexa Fluor 488 (Cat. A-11001), and Goat anti-Rat IgG (H+L) Alexa Fluor 568 (Cat. A-11077) at a 1:700 dilution for 1 hour at R.T. Finally, nuclei were counterstained with Hoechst 33342 (J62134.100, Thermo Fisher Scientific) for 10 minutes. Images were captured using a fluorescence microscope, and the expression of the markers was analyzed to assess the pluripotency of the iPSCs.

### Brain organoids Protocol and characterization

For the generation of brain organoids, we started with an 80%-90% confluency of human pluripotent stem cells (hPSCs) cultured in feeder-free conditions.

Cells were washed once with Dulbecco’s phosphate-buffered saline and dissociation was conducted using 1mL of Accutase solution (A6964, Merck). Following detachment, the cells were centrifuged and resuspended using mTeSR Plus media (100-0276, STEMCELL Technologies) containing Y-27632 dihydrochloride (1254/10, Tocris) and FGF2 (233-FB-025/CF, R&D Systems). Cells were counted using Trypan Blue and were seeded in a non-treated U-shape 96 well plate. Cells were allowed to rest and form cell-to-cell connections for 24 hours before replacing the media. The day after the seeding, the media was replaced and the cells were treated with Neurobasal Media (Gibco) containing GlutaMAX (Gibco), Penicillin-Streptomycin, N-2 (Gibco), B-27 (Gibco) and supplemented with dual SMAD inhibitors and vitamin A. After the formation of stable 3D spheres, each sphere is transferred into an ultra-low attachment flat 24-well plate for the continuation of neuronal differentiation and maturation with the same Neurobasal media supplemented with fibroblast growth factor 2 (basic), transforming growth factor beta 1, and gamma-secretase inhibitor.

Cryosectioning and immunostaining were performed to analyze the expression of specific markers in organoid samples. Organoids were harvested and fixed with 4% paraformaldehyde for 1 hour at room temperature (R.T.). Following fixation, the organoids were embedded in an optimal cutting temperature (OCT) compound and frozen at -80°C. Sections of 10 μm thickness were obtained using a cryostat and mounted on glass slides. The sections were permeabilized with 0.2% Triton X-100 for 30 minutes at R.T. and blocked with 5% bovine serum albumin (BSA) for 1 hour at R.T. After blocking, the sections were incubated overnight at 4°C with primary antibodies: Mouse anti-Sox2 (SC-365823, Santa Cruz Biotechnology) and Rabbit anti-Tuj1 (GTX129913-25, GeneTex) at concentrations recommended by the manufacturer. Sox2 staining was used as a marker for iPSCs, as it is a critical transcription factor essential for maintaining pluripotency and self-renewal in stem cells. Tuj1, a marker for beta-III tubulin, was used to identify differentiated neurons within the organoids, as Tuj1 is a well-established marker for neuronal differentiation. Following primary antibody incubation, the sections were washed with PBS and incubated with secondary antibodies conjugated to fluorophores, such as Goat anti-Mouse IgG (H+L) Alexa Fluor 488 and Goat anti-Mouse IgG (H+L) Alexa Fluor 568, for 2 hours at R.T. Finally, the sections were counterstained with Hoechst for 10 minutes to visualize the nuclei. Images were captured using a fluorescence microscope, and the expression of the markers was analyzed to assess the pluripotency of the iPSC-derived organoids and the extent of neuronal differentiation.

### Electrophysiological Recording and Stimulation System

Organoids were plated on multielectrode arrays (MEA, 60MEA200/30iR-Ti, Multichannel system), coated with PEI (Polyethyleneimine, 0.1%).

The RHS Stim/Recording Controller, connected to the RHS2116 stim/amplifier chips (Intan Technologies, Santa Monica, CA), was used for electrophysiological measurements. Broadband electrophysiological signals were acquired at a rate of 30 kHz, with a 50Hz notch filter enabled and Intan amplifier bandwidth set to 1.17 – 7.60 kHz. Spontaneous activity was recorded for 5 minutes to determine baseline activity. Then, three cycles of electrical stimulation were administered, each lasting 20 seconds and comprising 20 biphasic pulses (100 μA, phase width 66.7 μs, duration 10 ms) at a frequency of 100 Hz. Subsequently, data was collected for an additional five minutes to measure the stimulation effect.

### MEA Data Analysis

Raw electrophysiological data were extracted from Intan files using the Intan RHX software and processed in Python. The signals were organized into a structured dictionary for downstream analysis. Each detected event underwent spike sorting. Electrodes with fewer than or equal to 2 spikes per minute were considered inactive. For active electrodes, we quantified several metrics including mean firing rate, mean spike amplitude, burst activity (single-channel bursts [SCB], network bursts [NB]), as well as network connectivity.

The mean firing rate was calculated as the number of spikes per second. Spike amplitude was defined as the minimum voltage of the spike waveform after baseline correction. A burst was defined as at least five spikes occurring within a 30 ms window. A network burst was defined as single channel bursts in at least five electrodes within a 500 ms window. Network connectivity was assessed by comparing the observed number of spike co-occurrences to the expected number under random conditions, based on the mean firing rate in each electrode.

Dimensionality reduction was performed using Principal Component Analysis (PCA) implemented via the scikit-learn package. High-dimensional electrophysiological features were projected into a 3D space, enabling both visual and quantitative comparisons across experimental groups. In addition, radar plots were used to visualize group-level averages of standardized biological features, including firing rate, connectivity, and bursting dynamics, across all conditions.

### Data analysis

Data visualization and statistical analysis were performed using GraphPad Prism (GraphPad Software, San Diego, CA). Appropriate statistical tests – including t-tests, one-way ANOVA, Kruskal-Wallis H test, and relevant post-hoc analyses – were applied as indicated.

## Results

### From Urine to Patient-Derived Brain Organoids on MEA (Figure 1)

Urine samples were collected and centrifuged to isolate the epithelial cell fraction (Figure 1A). These cells were cultured and reprogrammed into induced pluripotent stem cell (iPSC) colonies (Figure 1B), which were validated using standard markers for cell viability (Hoechst) and pluripotency (NANOG) (Figure 1C). iPSCs were then differentiated into multiple patient-derived brain organoids (Figure 1D–E) and characterized by immunostaining for cell viability (Hoechst), neural progenitor markers (SOX2), and early neuronal maturation (TUJ1). Morphological organization of the organoids is shown at both the whole-organoid level (top panels) and at higher magnification, highlighting neural vesicle structures (bottom panels) (Figure 1G–J). The functional platform, comprising patient-derived brain organoids cultured on a 64-electrode multi-electrode array (MEA), is illustrated in Figure 1F.

**Figure 1:**
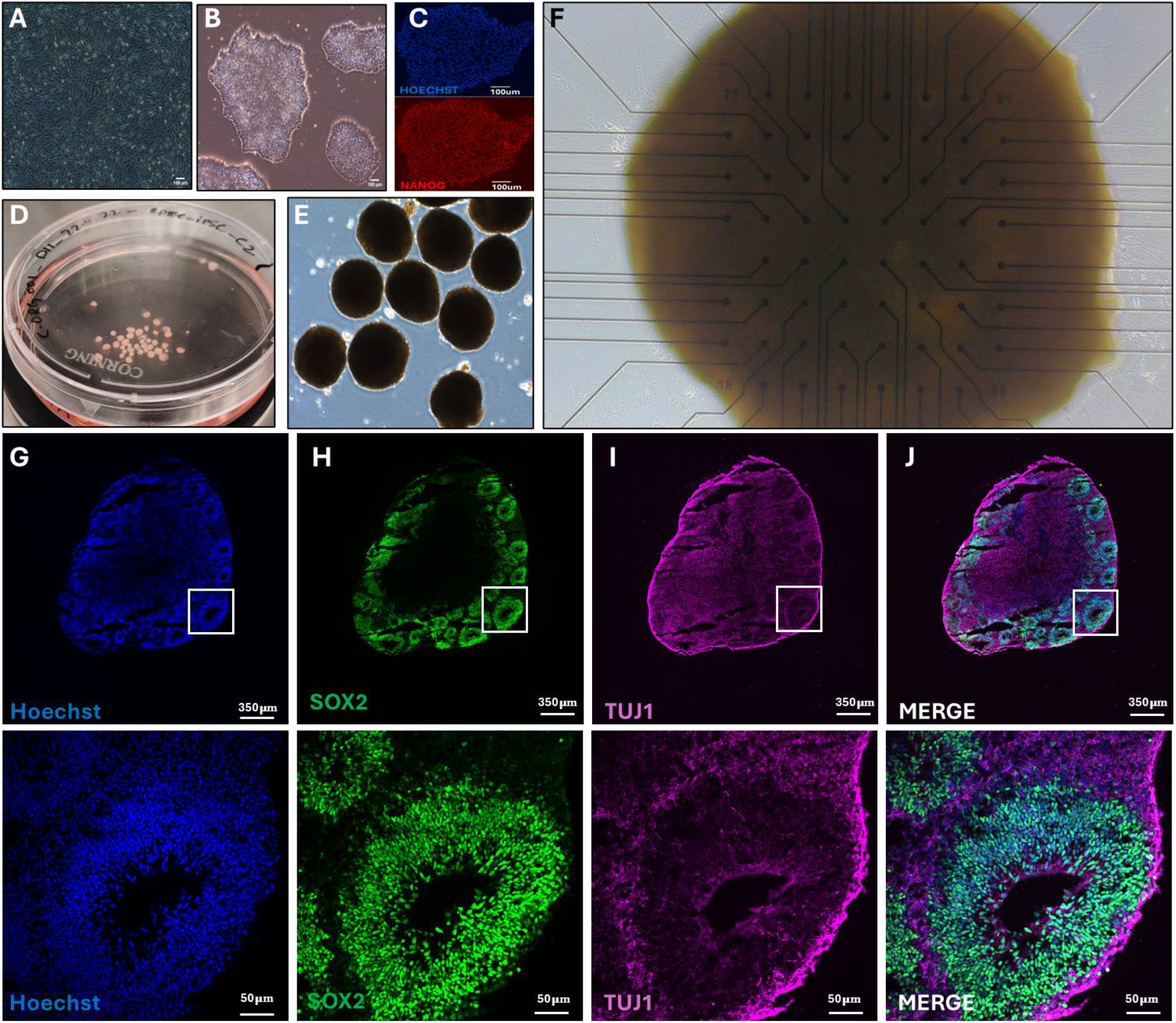
Urine epithelial Patient-derived brain organoids formation and characterization. **A**. Patient-derived epithelial progenitor. **B**. iPSCs colonies generation. **C**. iPSC characterization for viability (Hoechst) and pluripotency (NANOG). **D-E**. Visualization of multiple brain organoids formed from IPSC colonies. **F**. Multielectrode array recordings of patient-derived brain organoid set-up. **G-J**. Brain organoids characterization. G. Hoechst for vitality characterization, H. SOX2 for neuronal progenitor stem cells, I. TUJ1 for neuronal early maturation characterization, J. Merged photo: Top – whole organoid imaging; bottom -enlarged neural vesicle image.

### Characterization and Comparison of Basic (resting state) Electrophysiological Properties of Patient-Derived Brain Organoids from Syndromic, and Idiopathic ASD Patients, and Controls (Figure 2)

We characterized the spontaneous, resting-state electrophysiological activity of brain organoids derived from syndromic and idiopathic autism spectrum disorder (ASD) patients and compared them to neurotypical controls (Figure 2). Representative four-second recordings illustrate apparent differences in neuronal dynamics across groups, including patterns of hyperactivity, hypoactivity, altered burst synchrony, and amplitude variability. Each ASD subpopulation is consistently color-coded throughout the study for clarity (Figure 2B).

**Figure 2:**
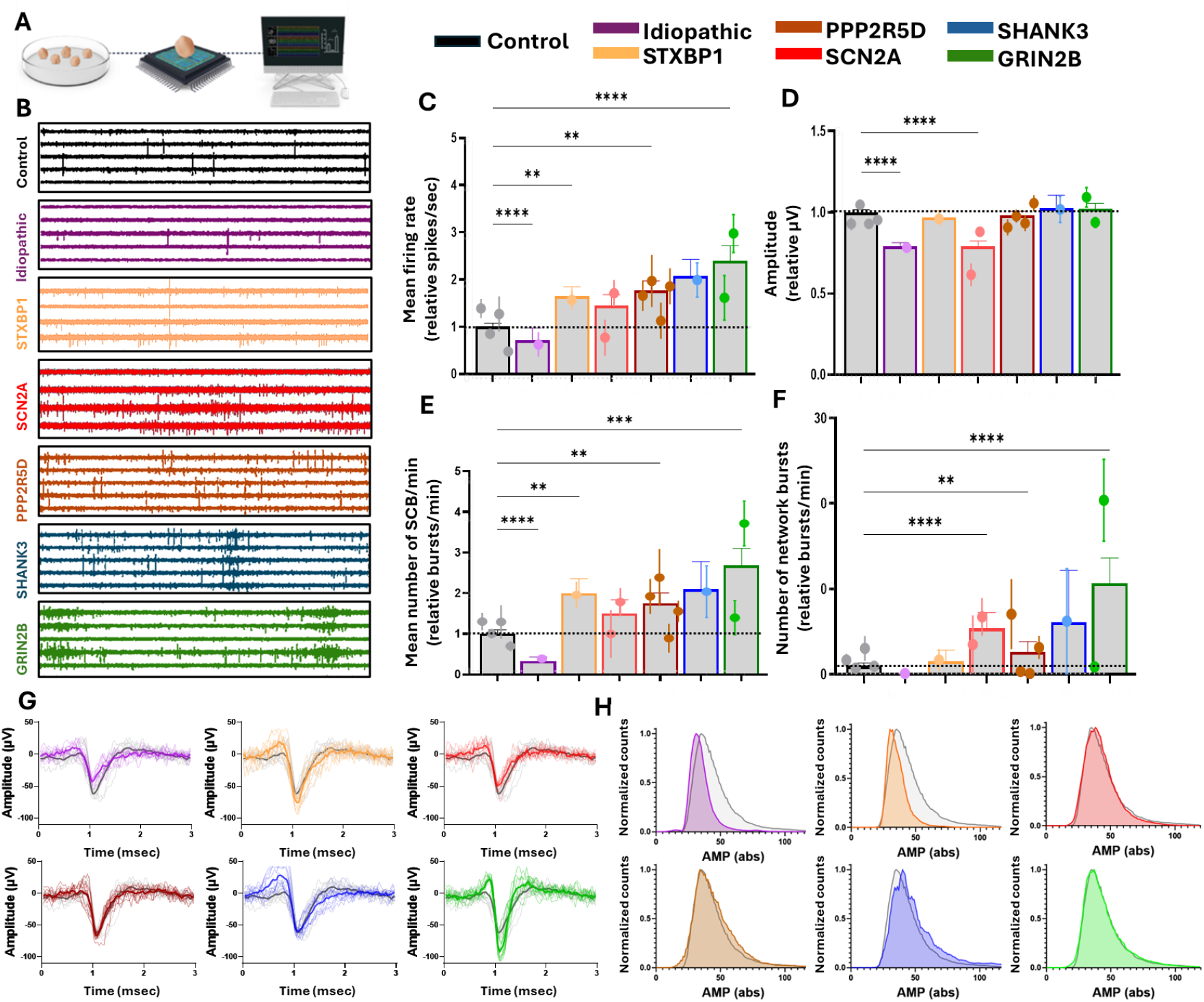
Characterization, comparison, and clustering of patient-derived brain organoids’ basic electrophysiological parameters (resting-state). **A**. Experimental setup graphic scheme. **B**. Representative recordings of electrophysiological activity of patient-derived brain organoids from different ASD subpopulations with their color coding. **C**. Mean firing rate (MFR) of different sub-populations represented relative to control. **D**. Amplitude of different sub-populations represented relative to control. **E**. Single channel burst (SCB) of different sub-populations represented relative to control **F**. Number of network bursts (NB) of different sub-populations represented relative to control. **G**. Spike shape of each ASD subpopulation compared to the representative spike of the control. **H**. Normalized amplitude histograms of each ASD sub-population compared to the control. (C-F, Dots are the average of all the brain organoids of each patient, and lines are SEM of all the brain organoids of each patient. One-way ANOVA (C-F), *>0.05, **>0.01, ****>0.0001, Scheffe).

We quantified several key electrophysiological features—mean firing rate, signal amplitude, and mean number of single-channel bursts. All ASD-derived organoids displayed significant deviations from the control group in at least one of these baseline parameters. The idiopathic ASD organoids showed a consistent hypoactive profile, with significantly reduced mean firing rate, amplitudes, and single-channel bursts compared to controls (all p < 0.0001).

In contrast, most syndromic ASD subtypes exhibited significantly increased mean firing rates, including STXBP1 (p < 0.01), PPP2R5D (Jordan’s syndrome) (p < 0.02), and GRIN2B (p < 0.0001). The SCN2A-derived organoids revealed heterogeneity in firing rate across patient lines, with a significant difference between them. Notably, both SCN2A lines showed significantly reduced amplitudes compared to controls (p < 0.0001), independent of their differing firing rates.

Amplitude dynamics are visualized in Figure 2G, where 20 randomly selected signals from each group are overlaid on the mean signal and the control average for comparison. The normalized amplitude distribution is presented in Figure 2H.

### Characterization of Short-Term Synaptic Plasticity in Patient-Derived Brain Organoids from Syndromic and Idiopathic ASD Patients and Neurotypical Controls (Figure 3)

To investigate short-term synaptic plasticity across brain organoids, we applied a three-cycle electrical stimulation paradigm using a multi-electrode array (MEA) to evoke short-term depression (STD) and short-term potentiation (STP) responses. The distribution of STD- and STP-dominant responses, as well as the unchanged portion, is illustrated in the pie charts (Figure 3B). The figure illustrates a predominantly STD response (blue) for all sub-populations, with STP (red) accounting for a minority of responses.

**Figure 3:**
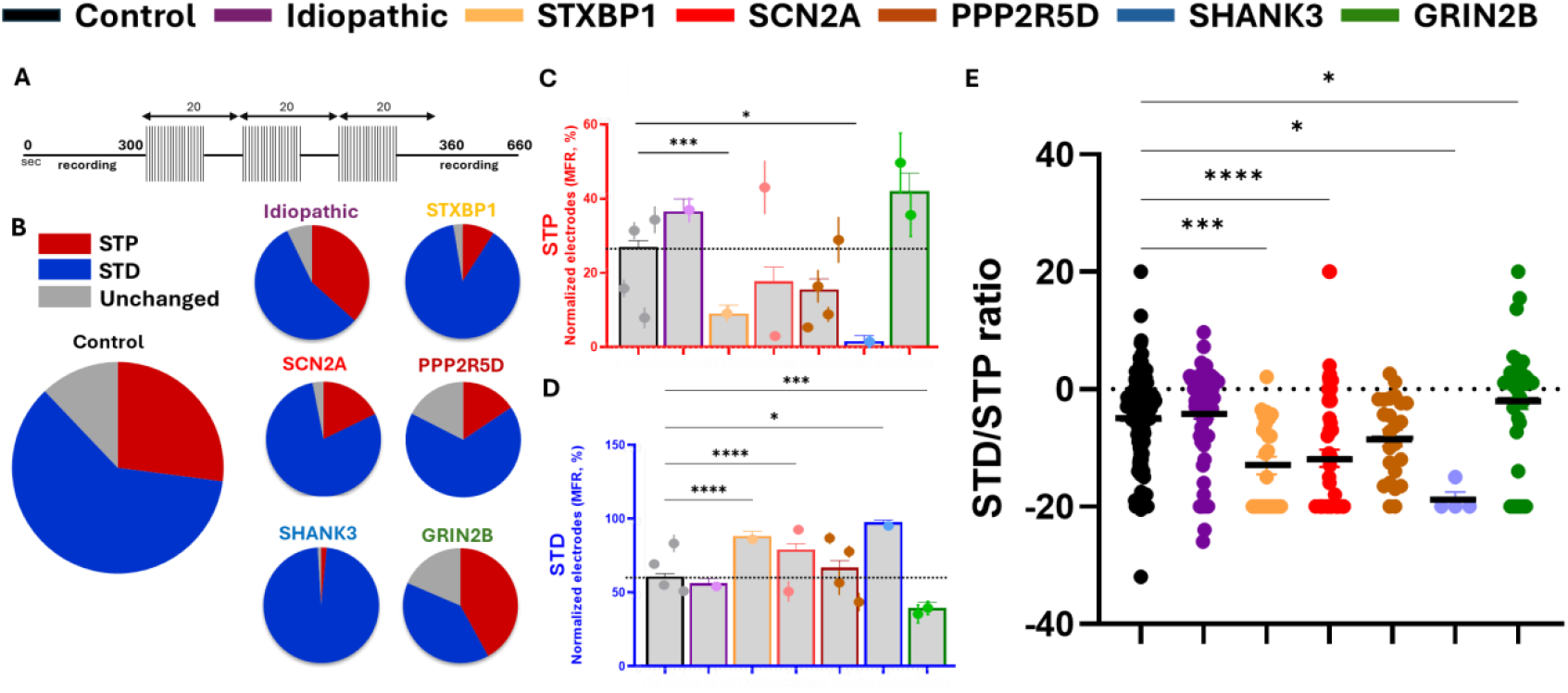
Characterization of short-term plasticity and comparison of various patient-derived brain organoid lines in ASD. **A**. Stimulation protocol scheme. **B**. Averages of STD, STP and unchanged electrodes presented in pie charts **C**. MFR-based STP in each sub-population of patient-derived brain organoids. **D**. MFR-based STD in each sub-population of patient-derived brain organoids. **E**. MFR-based STP/STD ratio in each sub-population of patient-derived brain organoids, with positive values representing a higher STP relative to STD (STP/STD), and negative values indicating a higher STD relative to STP (STD/STP). The MFR-based STP/STD ratio in each sub-population represents varying depressive tendency of brain organoids. (One-way ANOVA, *>0.05, **>0.01, ***>0.001, Scheffe).

Quantitative analysis of STP responses revealed significantly reduced STP response in several syndromic lines compared to controls (Figure 3C) including STXBP1 (p < 0.001), and SHANK3 (p < 0.05) lines as well as one of the SCN2A lines (p < 0.001). Complementary data from STD analysis revealed a significant increase in STD for these lines (Figure 3D). This mirrored response suggests the increase of the STD portion to some extent, occurred at the expense of the STP portion.

Notably, GRIN2B-derived organoids exhibited a slightly elevated STP response compared to the control lines (Figure 3C), while demonstrating a significant reduction in STD (p < 0.01, Figure 3D). This suggests that the reduction in STD was accompanied by a larger proportion of unchanged responses in the mean firing rate. In contrast to the aforementioned lines, no significant differences were observed in STP or STD amplitudes for the idiopathic and PPP2R5D-derived organoids relative to controls. These findings highlight the variations in STD and STP responses observed among ASD subpopulations, suggesting group-specific plasticity dynamics.

Lastly, individual data points of the STD/STP ratio for each sub-population are presented in Figure 3E, with positive values reflecting a higher STP relative to STD (STP/STD) and negative values indicating a higher STD relative to STP (STD/STP). The data demonstrates variability across patient-derived organoids with a similar average ratio observed for the control group and the idiopathic ASD line. The STXBP1, SCN2A, PPP2R5D, and SHANK3 groups exhibited a significant negative shift in the STD/STP ratio, whereas GRIN2B organoids presented a slight positive shift and displayed the most balanced STD/STP distribution, suggesting a deviation from typical short-term depression mechanisms.

Together, these results suggest that specific genetic mutations associated with syndromic ASD can significantly disrupt short-term synaptic plasticity. The observed differences in STD and STP across ASD subtypes support the hypothesis that synaptic dysfunction in ASD is highly heterogeneous and genotype-dependent.

### Network Connectivity Analysis and Comparison of Pre- and Post-Electrophysiological Stimulation in Patient-Derived Brain Organoids (Figure 4)

To evaluate network-level functional connectivity and its responsiveness to external stimulation across patient-derived brain organoids, we applied a Cross-Frequency Phase synchronization (CFP) analysis on spontaneous neuronal activity recorded in 30-second bins before and after electrical stimulation (Figure 4A). This method enabled quantification of dynamic changes in network structure, including network size and average connectivity strength, across both baseline and evoked states.

**Figure 4:**
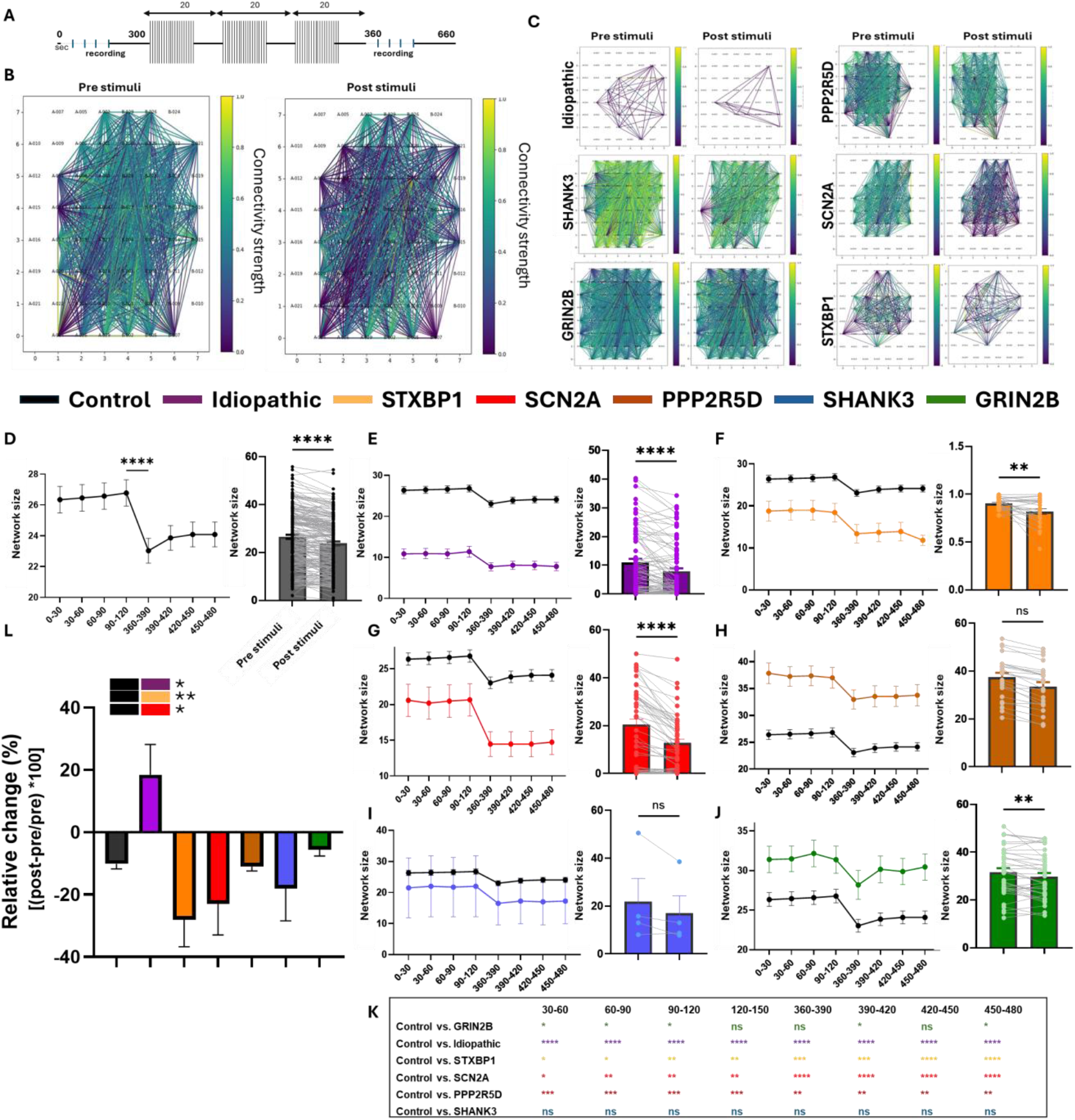
Network size and connectivity dynamic analysis and comparison between different ASD lines of patient-derived brain organoids. **A**. Stimulation protocol scheme, the data were divided into 30-second bins to follow the dynamics. **B**. Representative analysis of connectivity pre- and post-stimuli in the control group. **C**. Representative analysis of connectivity pre- and post-stimuli in ASD groups. **D**. Dynamic analysis of network size of the control group (left: time dynamics, right: pre- and post-stimuli comparison in paired analysis). **E-J**. Dynamic analysis of network size of the ASD groups, compared to control (black). **K**. Summary table of the statistical differences for each group compared to control at each time point. (Fisher’s exact test, *<0.05, **<0.001, ****<0.0001). **L**. The relative influence of the stimuli in percentages on the “network size” parameter. Statistics represent the comparison to the control group. (Kruskal-Wallis test, *<0.05, **<0.001, ****<0.0001)

Visual comparisons of network structure (Figure 4B–C) revealed noticeable differences between neurotypical controls and ASD-derived organoids. The control organoids exhibited relatively dense and stable connectivity patterns pre- and post-stimulation, while ASD-derived organoids showed greater variability, with some exhibiting fragmentation or hyperconnectivity, suggesting altered functional organization at the network level.

Temporal dynamics of network size were plotted over a 120-second window (Figure 4D–J). In the control group (Figure 4D), connectivity remained relatively stable pre-stimulation, followed by a modest and reproducible decline in network size post-stimulation. This pattern likely reflects an adaptive network response and reorganization following input. Paired t-test analysis confirmed a significant post-stimulation decrease in network size in the control group (p < 0.05).

In contrast, ASD-derived organoids displayed heterogeneous and often dysregulated responses to stimulation:

1. **Idiopathic ASD organoids** (Figure 4E) showed reduced adaptability, with minimal post-stimulation change in network size, diverging significantly from the control profile at multiple time points (p < 0.05).
2. **STXBP1-derived organoids** (Figure 4F) displayed a pronounced and early collapse in network size post-stimulation, with a significant paired reduction (p < 0.001), suggesting impaired recovery and network fragility.
3. **SCN2A lines** (Figure 4G) also showed a significant network reduction post-stimulation (p < 0.01), though with a delayed onset compared to STXBP1.
4. **PPP2R5D (Jordan’s syndrome) organoids** (Figure 4H) maintained higher pre-stimulation network size but demonstrated a sharp drop immediately following stimulation (p < 0.0001), indicating potential hyperconnectivity at baseline with impaired dynamic regulation.
5. **GRIN2B organoids** (Figure 4I) showed the most consistent deviation from control, with an attenuated and erratic response across the full time window (p < 0.05 at multiple time points).
6. **SHANK3 organoids** (Figure 4J) showed a moderate reduction in connectivity, statistically distinct from the control profile at late time points (p < 0.05).

Statistical comparisons at each time point between ASD subgroups and the control (Figure 4K) confirmed significant deviations in connectivity dynamics in multiple ASD lines, particularly STXBP1, GRIN2B, and PPP2R5D.

Finally, Figure 4L summarizes the relative change in connectivity, calculated as a percentage [(post-pre)/pre × 100], offering a quantitative measure of the stimulus-driven shift in network activity. STXBP1, SCN2A, PPP2R5D, and GRIN2B organoids all demonstrated significantly greater relative reductions in network size compared to controls, while SHANK3 and idiopathic ASD organoids showed more subtle or variable shifts.

Together, these results indicate that patient-derived organoids from different ASD subtypes exhibit distinct network-level phenotypes, with altered functional connectivity dynamics and reduced plasticity in response to stimulation. These patterns appear to be mutation-specific, supporting the use of network-level analysis as a discriminative biomarker of circuit dysfunction in ASD.

### Multidimensional Characterization and Comparison of Electrophysiological Features Within and Between Groups (Figure 5)

To summarize and visualize the diversity in electrophysiological features across all samples, we applied principal component analysis (PCA), which enabled us to reduce 18 electrophysiological parameters into a three-dimensional space based on their similarity (Figure 5A). In this clustering space, each patient is represented by a dot (representing the mean of their brain organoid data) and error bars (representing the standard error of the mean, or SEM), allowing for the assessment of both intra-subject and inter-subject variability. This approach revealed that variation between organoids from the same individual was relatively small, while inter-subject differences were more significant, especially between ASD patients and controls. The control group clustered closely, indicating minimal variability, whereas both syndromic and idiopathic ASD samples showed broader dispersion.

**Figure 5:**
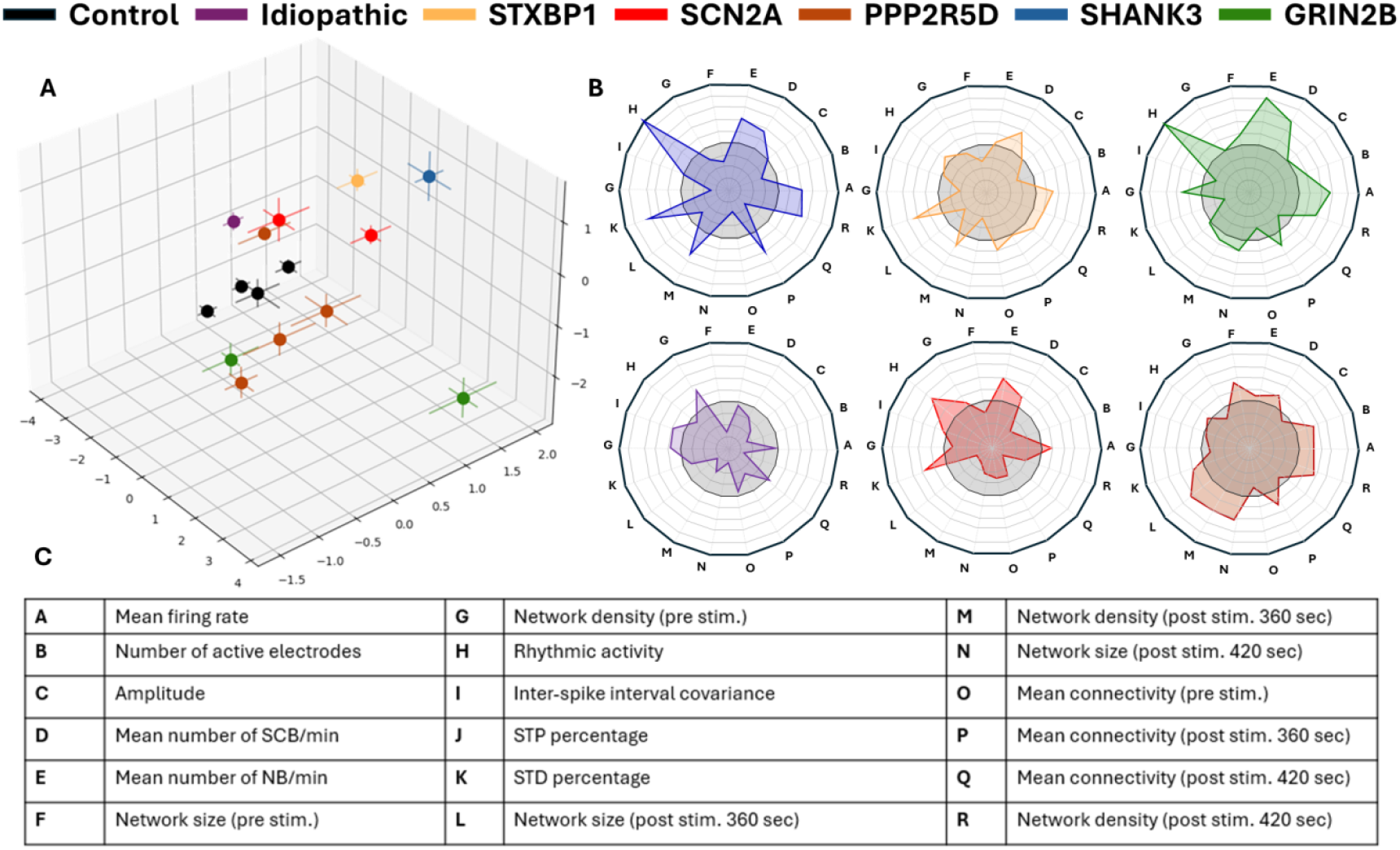
Multidimensional clustering of electrophysiological parameters in brain organoids from ASD subpopulations. **A**. Principal component analysis combines 18 parameters into 3D dimensions. Each patient’s data is represented by the mean (dot) and the standard error of the mean (lines) from all the brain organoids of that patient. **B**. Radar plots show the impact of various parameters, compared to the control (normalized to 1, gray rectangle). **C**. List of the features included in the PCA and radar plot analyses.

To better understand which electrophysiological parameters contributed to group differentiation, we used radar plots normalized to the control group (value = 1), highlighting deviations in ASD groups across multiple features (Figure 5B). A complete list of the 18 parameters included in the PCA is provided in the accompanying table (Figure 5C).

## Discussion

In this study, we aimed to extensively characterize and compare the electrophysiological patterns of brain organoids generated from individuals with syndromic and non-syndromic ASD, including cases with **SHANK3 haploinsufficiency** (Phelan-McDermid syndrome), **PPP2R5D** (Jordan’s syndrome), **GRIN2B syndrome, SCN2A syndrome**, and **STXBP1 syndrome**, relative to healthy controls. While previous studies have reported molecular and electrophysiological alterations in ASD brain organoids, including in SHANK3 haploinsufficiency^26^, our study provides a novel and systematic comparison across multiple ASD subtypes. We examined organoids derived from individuals with both syndromic and non-syndromic autism and found distinct electrophysiological alterations relative to controls. These abnormalities affect core aspects of neuronal coding—an essential feature of functional neural networks and cognitive processing. Here, we analyzed over 400 brain organoids derived from eleven individuals with diverse autism etiologies and four neurotypical controls. We identified distinct electrophysiological properties both between individuals with autism and controls, and across different monogenic forms of autism. Both resting-state electrophysiological properties and evoked responses (short-term potentiation and short-term depression) were studied, along with network connectivity analysis. Since electrophysiological data contains a high number of parameters, each of which can be influenced differently from the biological basis of the neurons and the organism, we used dimensionality reduction and a machine learning algorithm (PCA) to standardize the data for coherent comparison. The results, represented by the mean and standard error of the mean (SEM) of all brain organoids from each patient, highlighted the relative resemblance of the brain organoids from the control patients against the high variability among the different ASD patients.

It has been widely reported that syndromic patients can still present with different clinical phenotypes, even when carrying the same genetic mutation^27–29^. This clinical heterogeneity can be attributed not only to gene–environment interactions but also to inherent biological variation at multiple regulatory levels. Factors such as non-coding regulatory elements (e.g., enhancers, silencers, and long non-coding RNAs) can differentially modulate gene expression among individuals, despite identical coding mutations. Furthermore, variations in post-translational cellular processes including protein folding, trafficking, phosphorylation, and ubiquitination— can result in diverse functional outcomes from the same genetic alteration. These multilayered regulatory mechanisms contribute to the complexity of genotype–phenotype relationships and offer a compelling explanation for inter-individual variability in clinical presentation.

Consistent with this, we found that patients diagnosed with the same ASD syndrome (e.g., GRIN2B-related syndrome) exhibited distinct patterns of electrical activity and were not always clustered into the same group. This divergence in electrophysiological profiles aligns with the clinical differences observed between these patients (Table S1), reinforcing the notion that shared genetic mutations do not necessarily lead to uniform phenotypic outcomes. For example, only one of the GRIN2B lines has reported seizures, which was also manifested, to some extent, in an abnormal and unique rhythmic burst activity. This rhythmic burst activity was observed in some of the patients who had seizures (Figure S3). This emphasizes the importance of functional electrophysiological data as additional clinical diagnostic information on the subject, in addition to the syndromic and genetic characterizations, for a better understanding of the heterogeneity within the same syndromic ASD and between different ones. Additionally, since the vast majority of ASD cases are idiopathic and non-syndromic, the electrophysiological information may be used for better clustering and characterization of sub-populations within the non-syndromic population.

Furthermore, while animal models have been used for several decades to study neurological and psychiatric disorders, their translational relevance remains limited, mainly in drug testing and development, with less than 8% success in translation from pre-clinical studies to clinical practice^30^. This challenge is further compounded by the human brain’s biological complexity, making it difficult to develop reliable animal models even for monogenic neurodevelopmental disorders. Despite a known single-gene mutation, these models often fail to accurately recapitulate the human phenotype. This limitation is even more pronounced in complex multifactorial conditions such as ASD, where the interplay of genetic and environmental factors cannot be easily replicated in animal systems.

In summary, we provide a comprehensive electrophysiological characterization of brain organoids derived from individuals with both syndromic and non-syndromic ASD, compared to neurotypical controls. By analyzing over 400 organoids from diverse ASD etiologies, we identified distinct patterns of neuronal activity across and within ASD subtypes, including heterogeneity among individuals with the same genetic mutation. These alterations, observed in resting-state activity, evoked responses, and network dynamics, were further resolved using dimensionality reduction. Notably, our study highlights the critical value of functional electrophysiological readouts in revealing ASD-specific signatures and distinguishing subpopulations, even within genetically defined syndromes. This underscores the potential of patient-derived brain organoids to serve not only as a complement to genetic diagnosis but also as a powerful translational platform for uncovering mechanistic insights into ASD heterogeneity, enabling diagnostics in non-syndromic or idiopathic cases, and guiding future personalized therapeutic strategies^31–34^.

## Acknowledgments

We are grateful to all of the families at the participating Simons Searchlight sites as well as the Simons Searchlight Consortium, formerly the Simons VIP Consortium. We appreciate obtaining access to biospecimens, and phenotypic data on SFARI Base. Approved researchers can obtain the Simons Searchlight population dataset described in this study by applying at https://base.sfari.org

## Conflict of Interest Statement

Itay&Beyond is a for-profit company established to develop novel therapeutics and discovery tools for neuropsychiatric disorders. Prof. Tenenbaum serves on the advisory board of Itay&Beyond.

## Supplementary Information

**Table S1:** Clinical summary of this study’s patients and volunteer participants.

| Sample | Age [years] | Sex | Protein change | Genetic mutation | Co-Morbidity | Psychiatric | Medication |
| --- | --- | --- | --- | --- | --- | --- | --- |
| Control | 45 | M | - | - | - | - | - |
| Control | 35 | F | - | - | - | - | - |
| Control | <1 | Unknown | - | - | - | - | - |
| Control | 51 | F | - | - | - | - | - |
| Shank3 | 11 | F | Haploinsufficiency | InsG3680<sup>(+/-)</sup> | Infection, seizures, developmental delay | - | - |
| Idiopathic | 13 | M | - | - | - | OCD | Yes |
| PPP2R5D 15653-x1 | 9 | M | Glu198Lys | 592G>A | Allergy, short stature, failure thrive, diarrhea, GERD, infection, end stage renal disease, crossed eyes, astigmatism, nearsighted, developmental delay, clumsy, cranial nerve disorder, low and high muscle tone, large head size, spastic cerebral palsy, movement disorder, cerebral palsy | - | - |
| PPP2R5D 16020-x1 | 15.5 | F | Glu420Lys | 1258G>A | Constipation, large head size, movement disorder, low muscle tone, seizures, crossed eyes | - | - |
| PPP2R5D 15655-x1 | 2 | F | Glu198Lys | 592G>A | GERD, large head size, cort blind, low muscle tone, repetitive eye movements, crossed eyes, astigmatism, sensory integration disorder, seizures | - | Yes |
| PPP2R5D 16051-x1 | 14.5 | F | Glu200Lys | 598G>A | Allergy, irregular menses, hemangioma, developmental delay, seizures in sleep, constipation, hypertension, infection, clumsy, large head size, low muscle tone | Attention deficit hyperactivity disorder | Yes |
| GRIN2B 15413-x1 | 9.5 | F | Gly611Val | 1832G>T | Allergy, hemangioma, short stature, constipation, GERD, infections, clumsy, small head size, low muscle tone, seizures, crosses eyes | - | Yes |
| GRIN2B 15265-x1 | 5.5 | M | Gly820Ala | 2459G>C | Failure thrive, constipation, GERD, infection, low muscle tone, respiratory, osteoporosis, crosses eyes | - | - |
| SCN2A 15473-x1 | 4 | M | Glu1155Alafs*2 | 3464_3468 delAACAG | Allergy, constipation, clumsy, movement disorder, low muscle tone, seizures | Attention deficit hyperactivity disorder | Yes |
| SCN2A 15912-x1 | 1 | M | Val1601Leu | 4801G>T | Crossed eyes, nearsighted, seizures, cort blind, movement disorder, low and high muscle tone, hydronephrosis, infections, constipation | - | Yes |
| STXBP1 15936-x1 | 4 | F | Arg122* | 364C>T | Allergy, seizures, constipation, clumsy, movement disorder, low muscle tone | - | Yes |

**Figure S1:**
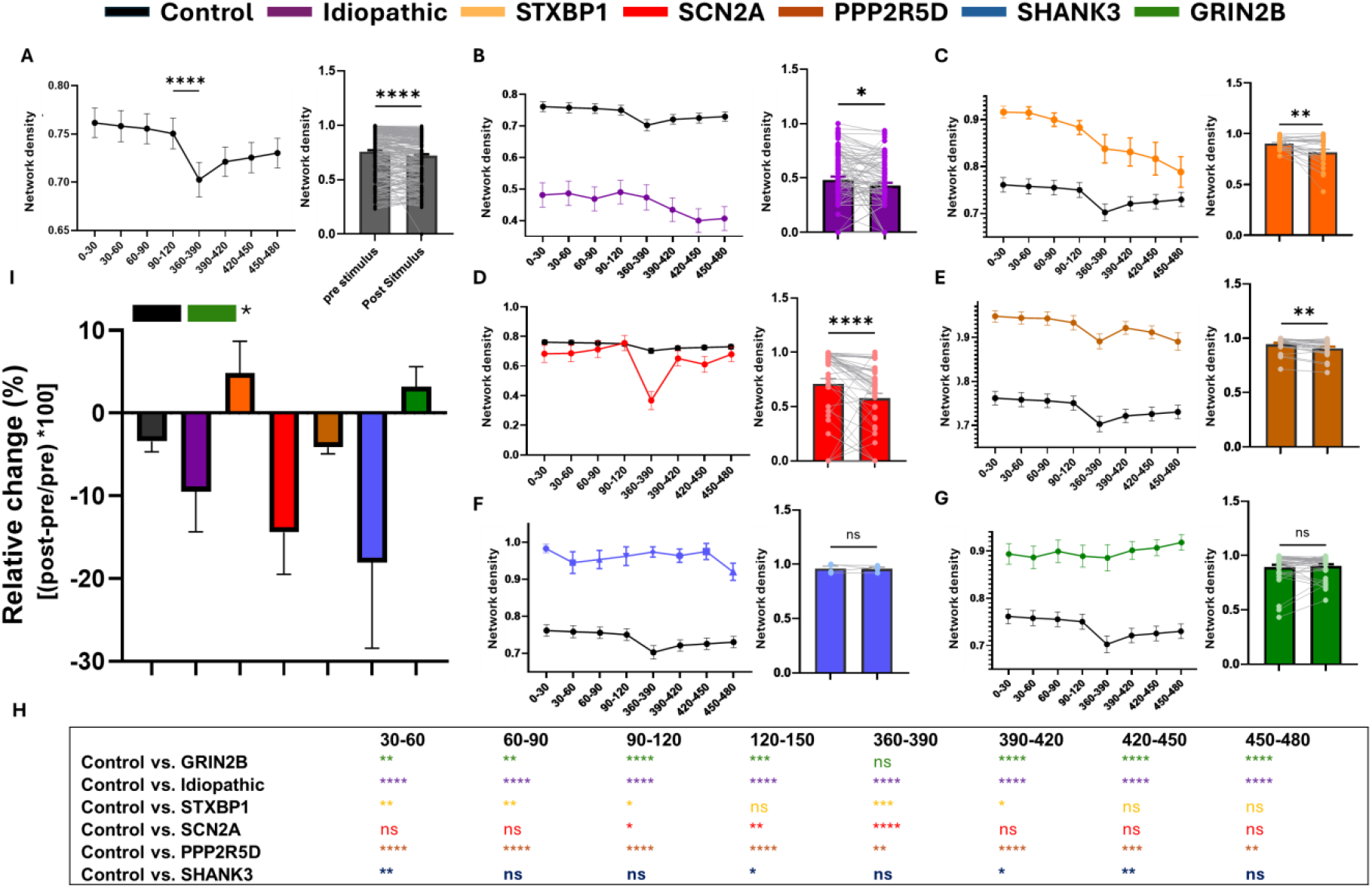
Network Density and connectivity dynamic analysis and comparison between different ASD lines of patient-derived brain organoids. **A**. Dynamic analysis of network density of the control group (left: time dynamics, right: pre- and post-stimuli comparison in paired analysis). **B-G**. Dynamic analysis of network size of the ASD groups, compared to control (black). **H**. Summary table of the statistical differences for each group compared to control at each time point. (Fisher’s exact test, *<0.05, **<0.001, ****<0.0001). **I**. The relative influence of the stimuli in percentages on the “network density” parameter. Statistics represent the comparison to the control group. (Kruskal-Wallis test, *<0.05, **<0.001, ****<0.0001).

**Figure S2:**
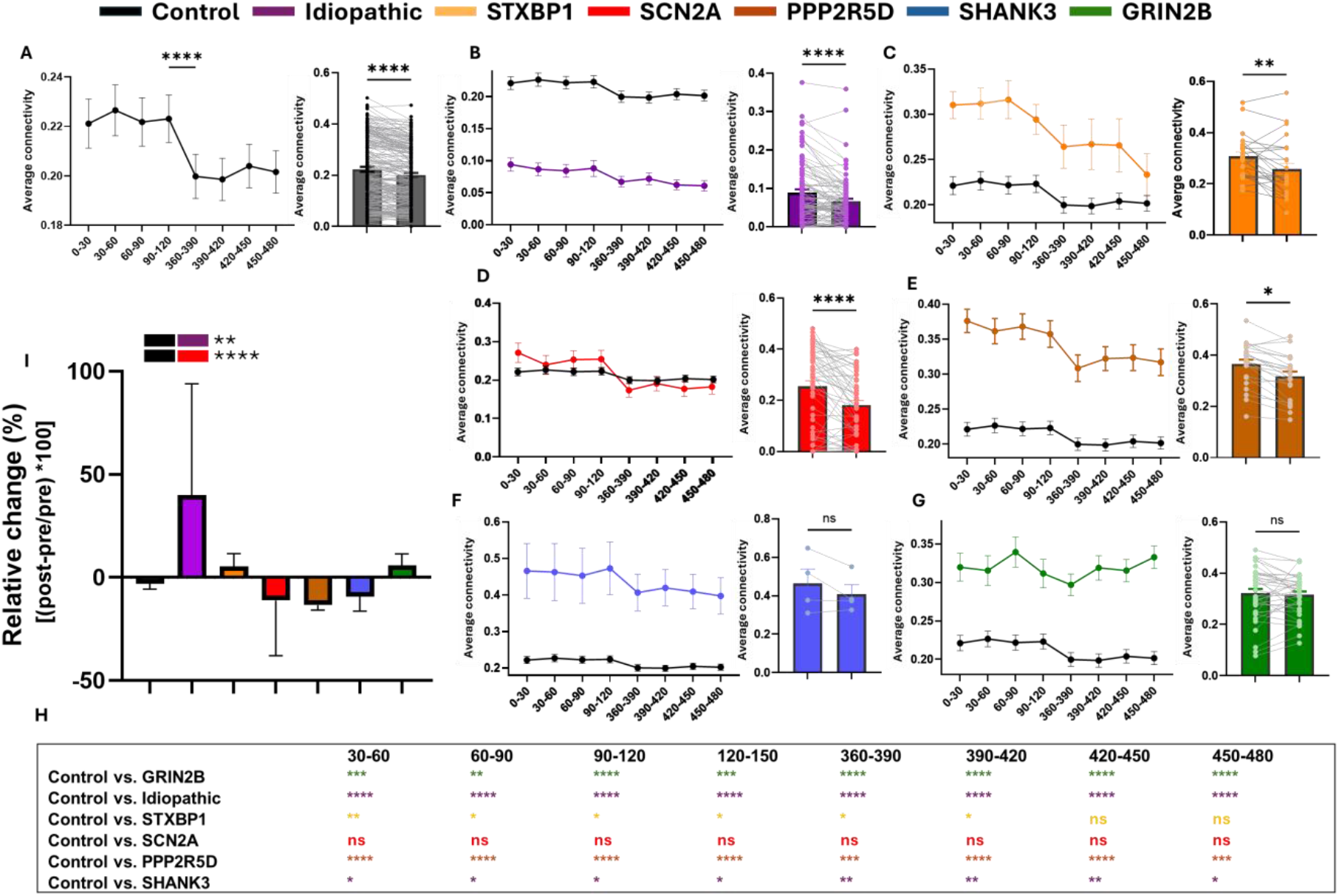
Network average connectivity and connectivity dynamic analysis and comparison between different ASD lines of patient-derived brain organoids. **A**. Dynamic analysis of average connectivity of the control group (left: time dynamics, right: pre- and post-stimuli comparison in paired analysis). **B-G**. Dynamic analysis of network size of the ASD groups, compared to control (black). **H**. Summary table of the statistical differences for each group compared to control at each time point. (Fisher’s exact test, *<0.05, **<0.001, ****<0.0001). **I**. The relative influence of the stimuli in percentages on the “average connectivity” parameter. Statistics represent the comparison to the control group. (Kruskal-Wallis test, *<0.05, **<0.001, ****<0.0001).

**Figure S3:**
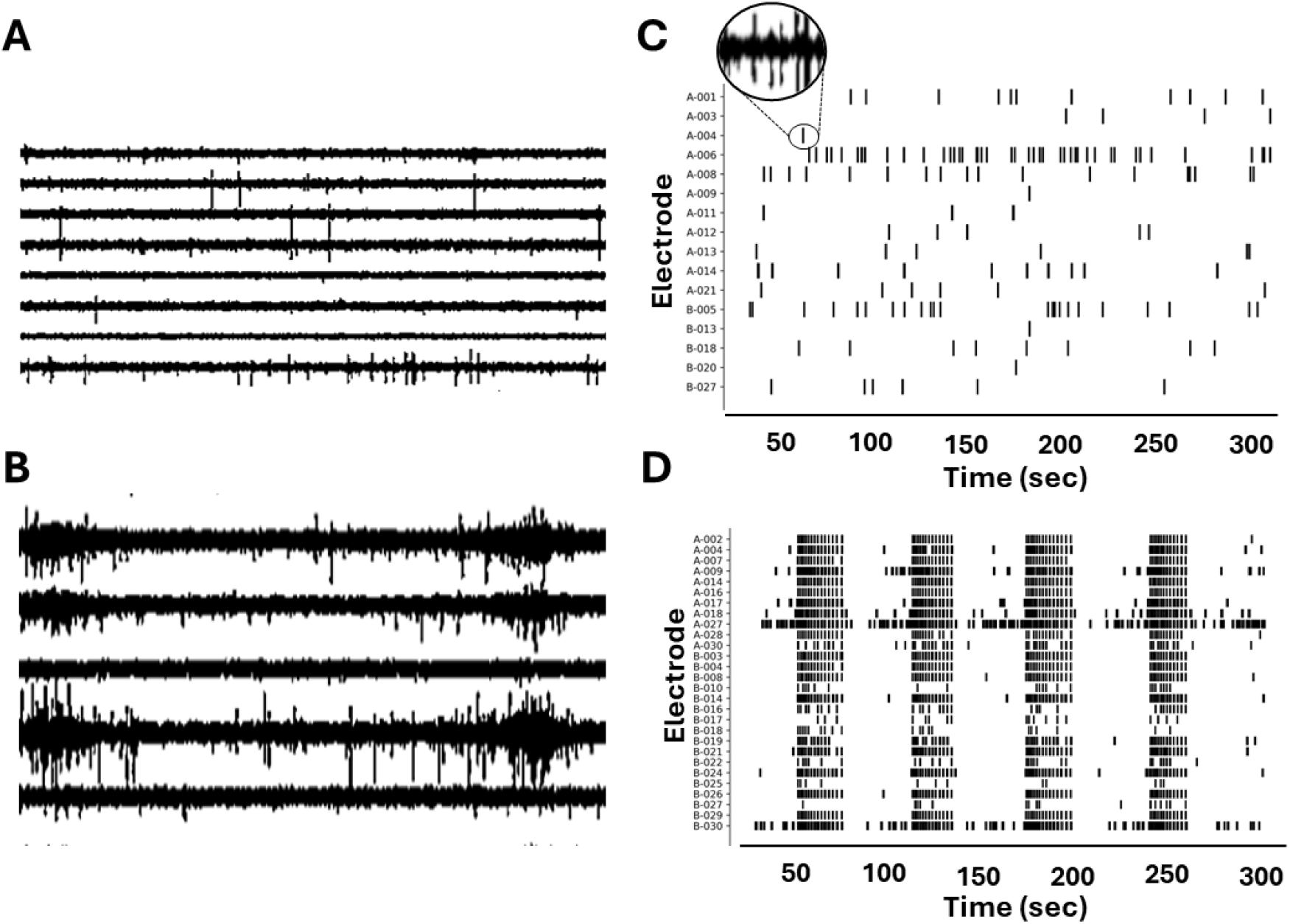
Abnormal rhythmic activity of synchronized bursts, presented by some of the patients with reported clinical seizures. **A**. Representation of the electrophysiological trace of a control patient. **B**. Representation of the electrophysiological trace of a patient with clinical seizures (GRIN2 B) **C**. Raster plot of the recordings from a control patient, each line is a single channel burst. **D**. Raster plot of the recordings from a patient with clinical seizures (GRIN2B), with each line representing a single-channel burst.

